# Phylogenetic reconstruction of ancestral ecological networks through time for pierid butterflies and their host plants

**DOI:** 10.1101/2021.02.04.429735

**Authors:** Mariana P Braga, Niklas Janz, Sören Nylin, Fredrik Ronquist, Michael J Landis

## Abstract

The study of herbivorous insects underpins much of the theory that concerns the evolution of species interactions. In particular, Pieridae butterflies and their host plants have served as a model system for studying evolutionary arms-races. To learn more about the coevolution of these two clades, we reconstructed ancestral ecological networks using stochastic mappings that were generated by a phylogenetic model of host-repertoire evolution. We then measured if, when, and how two ecologically important structural features of the ancestral networks (modularity and nestedness) evolved over time. Our study shows that as pierids gained new hosts and formed new modules, a subset of them retained or recolonized the ancestral host(s), preserving connectivity to the original modules. Together, host-range expansions and recolonizations promoted a phase transition in network structure. Our results demonstrate the power of combining network analysis with Bayesian inference of host-repertoire evolution to understand changes in complex species interactions over time.

---

For more than a century, biologists have studied the coevolutionary dynamics that result from intimate ecological interactions among species (Darwin 1877; Ehrlich and Raven 1964; Forister et al. 2012; Vienne et al. 2013). Butterflies and their host-plants are among the most studied of such systems; hence, various aspects of butterfly-plant coevolution have inspired theoretical frameworks that elucidate how interactions evolve in nature (Janz 2011; Suchan and Alvarez 2015). Two prominent conceptual hypotheses that explain host-associated diversification derive from empirical work in butterfly-plant systems: the escape-and-radiate hypothesis (Ehrlich and Raven 1964) and the oscillation hypothesis (Janz and Nylin 2008). The escape-and-radiate model predicts that butterflies (and host-plant lineages) diversified in bursts, resulting from the competitive release that follows the colonization of a new host (Futuyma and Agrawal 2009). Under this scenario, butterfly diversification should often be associated with complete host shifts, i.e. new hosts replace ancestral hosts (Fordyce 2010). In contrast, the oscillation hypothesis assumes that butterflies colonizing new hosts may retain the ability to use the ancestral host or hosts. According to this hypothesis, at any point in time, butterflies may be able to use more hosts than they actually feed on in nature. Defining the set of hosts used by a butterfly as its *host repertoire*, the oscillation hypothesis allows for a lineage to possess a realized host repertoire (analogous to realized niche) that is a subset of its fundamental host repertoire (Nylin et al. 2018). And while the fundamental host repertoire is phylogenetically conserved, the realized repertoire is less stable over evolutionary time, resulting in oscillations in the number of hosts used (i.e. host range). These oscillations in the realized host repertoire are thought to spur diversification.

Studies of various biological systems have supported the idea that distinct coevolutionary dynamics, such as those above, generate networks of ecological interactions with equivalently distinctive structural properties, such as modularity (Olesen et al. 2007) and nestedness (Bascompte et al. 2003). For example, nestedness has been associated with arms-race dynamics in bacteria-phage networks (Fortuna et al.2019). Other studies, however, suggest that some properties displayed by ecological networks are simply byproducts of the way these networks grow, e.g. by speciation-divergence mechanisms (Valverde et al. 2018, and references therein). Simulating data under a mixture of scenarios inspired by the escape-and-radiate and the oscillation hypotheses, Braga et al. (2018) suggested that nested and modular structures in extant butterfly-plant networks could be largely explained by a combination of the two scenarios. As a simulation-based framework, however, the Braga et al. (2018) approach cannot reconstruct how ancestral networks of ecological interactions evolved over deep time from biological datasets. Generally speaking, historical reconstructions of this kind would help to quantify when, under what conditions, and to what extent alternative scenarios of coevolutionary dynamics unfolded.

Beyond network analyses of extant species interactions, historical event-based inference methods are needed to identify the coevolutionary mechanisms that generated the observed interaction patterns. Even though ancestral ecological networks have been reconstructed using paleontological data (Dunne et al. 2014; Blanco et al. 2021), fossil-only approaches are not feasible for most groups of interacting species and over most geographical scales; this is the case with butterflies (Sohn et al. 2015) and butterfly herbivory (Opler 1973). Phylogenetic models offer an alternative method for inferring historical interactions. In the case of host-parasite coevolution, methodological constraints have hindered explicit modeling of host-repertoire evolution without significantly reducing the inherent complexity of the system. A newer phylogenetic method for modeling how discrete traits evolve along lineages (Landis et al. 2013) was recently extended to model host-repertoire evolution (Braga et al. 2020). Unlike previous approaches used to reconstruct past ecological interactions (e.g. Ferrer-Paris et al. 2013; Tsang et al. 2014; Jurado-Rivera and Petitpierre 2015; Navaud et al. 2018), the method of Braga et al. (2020) allows parasites to have multiple simultaneous hosts and to preferentially colonize new hosts that are phylogenetically similar to other hosts in each parasite’s host repertoire. These features reveal the entire distribution of ancestral host ranges at any given point in time, including the “long tail” of generalists often found in host-range distributions of extant species (Forister et al. 2015; Nylin et al. 2018), as well as temporal changes in host range. While the Braga et al. (2020) method was used to reconstruct plant-use characters for ancestral Nymphalini species, the study made no attempt to reconstruct ancestral ecological networks or network structures from its inferences.

For the present article, we performed a Bayesian analysis of host-repertoire evolution to reconstruct ecological networks on deep timescales from biological data. Here, we provide the means to test ideas about evolution of ecological networks by further developing the tool-set for analysis of inferred ancestral interactions. We show its usefulness by reconstructing ancestral Pieridae-angiosperm networks with a new probabilistic representation that makes fuller use of the posterior distribution of ancestral states. By reconstructing ancestral networks, we show how specific host shifts, host-range expansions, and recolonizations of ancestral hosts have shaped the Pieridae-angiosperm network over time, creating an evolutionarily stable modular and nested structure.

## Methods

### Pierid Butterflies and Angiosperm Hosts

We compiled an ecological interaction matrix from records of larval herbivory in butterfly genera by critically assessing previously published data (see Supplementary Information). To model how butterfly-plant interactions evolve, we used previously published time-calibrated phylogenies for 66 Pieridae genera (Edger et al. 2015, Fig. S1) and angiosperm families (Edger et al. 2015; Magallón et al. 2015). We pruned the host angiosperm phylogeny, keeping all 33 angiosperm families that are known to be hosts of pierid butterflies, then collapsing increasingly ancestral nodes until only 50 terminal branches were left. This increased the chance that all angiosperm lineages that ancestral butterflies might have once used as hosts were represented, while keeping the dataset manageable in size, as discussed in Braga et al. (2020).

### Model of host-repertoire evolution

We reconstructed how ecological interactions between Pieridae butterflies and their host plants evolved using a Bayesian phylogenetic modeling framework (Braga et al. 2020). The model treats the host repertoire as an evolving binary vector that indicates which host plants a particular butterfly lineage uses: host (1) or non-host (0). Each butterfly’s host repertoire then evolves along the branches of the butterfly phylogeny as a continuous-time Markov chain (CTMC) with relative rates of host gain (λ_1_) and loss (λ_0_) that are scaled by a global event rate (*μ*). In addition, the model includes a phylogenetic distance parameter (*β*) that, when *β* > 0, increases the relative rate of host gain for adding hosts that are phylogenetically similar to hosts currently being used by the butterfly. Following Braga et al. (2020), we measured phylogenetic distance between host lineages in two different ways: (1) using what we call the *anagenetic tree*, where distances reflect time-calibrated divergence times among hosts, and (2) using a modified *cladogenetic* tree, where all host branch lengths were set to 1, resulting in phylogenetic distances that are proportional to the number of older (i.e. family-level) cladogenetic events that separate two taxa. In this sense, the cladogenetic tree is equivalent to the kappa-transformed anagenetic tree with *κ* = 0 (Pagel 1994). We did not consider uncertainty in the host or butterfly phylogenies to facilitate the inference of model parameters under our data augmentation method, which may artificially reduce the uncertainty in our ancestral host repertoire estimates and any downstream analyses dependent on them. We also note that Braga et al. (2020) used three states (non-host, potential host and actual host) to model host repertoires, whereas we used only two because our data set did not report information on potential hosts. Refer to Braga et al. (2020) for further details about the model.

### Summarizing ecological interactions through time

Ancestral interactions were estimated by sampling histories of host-repertoire evolution during the Bayesian Markov chain Monte Carlo (MCMC) analysis (described below), meaning interaction histories were sampled jointly with model parameters from the posterior. We first summarized the sampled histories using a traditional representation of ancestral states (e.g. Nylin et al. 2014) by calculating the marginal posterior probabilities for interactions between each host plant and each internal node in the Pieridae phylogeny. Interactions with marginal posterior probability >0.9 were treated as ‘true’ occurrences, with all other interactions being treated as ‘false’. This traditional approach has three important limitations: (1) it only considers states at internal nodes, ignoring what happens along the branches of the butterfly tree; (2) by focusing on the highest-probability butterfly-plant interactions, it filters out ancestral interactions with middling probabilities; and (3) it is blind to how joint sets of interactions might have evolved together, as it is based on marginal probabilities of pairwise butterfly-plant interactions. We discuss each of these three aspects in detail below and explore new ways to summarize host-repertoire evolution.

#### Viewing ecological histories as networks

To resolve the first limitation, we reconstructed the host repertoires of all extant butterfly lineages at eight time slices, from 80 Ma to 10 Ma. Thus, instead of reconstructing the host repertoire of internal nodes in the butterfly tree, we reconstructed ancestral Pieridae-host plant networks at different ages throughout the diversification of Pieridae. This method captures more information about the system at specific time slices and, most importantly, can quantify changes in network structure over time, as contrasting hypotheses of eco-evolutionary dynamics are expected to generate similarly contrasting structures in ecological networks (Braga et al. 2018).

#### Summarizing posterior distributions of networks with point estimates

To investigate how much information is lost when we only consider the highest-posterior interactions (limitation 2), we compared three kinds of summary networks for each time slice: (1) binary (presence/absence) networks with probability thresholds of 0.9, (2) weighted networks with thresholds of 0.5, and (3) weighted networks with thresholds of 0.1. A binary network treats interactions with >0.9 marginal posterior probability as present, while all other interactions are considered absent. In the two remaining weighted network types, plant-butterfly interactions have weights equal to their posterior probabilities, but interactions with probabilities under a given threshold are assigned the weight of 0 (absent). The two weighted network types (numbered 2 and 3, above) exclude all interactions with probability <0.5 or probability <0.1, respectively.

To characterize the structure of extant and ancestral (inferred) networks, we used two standard metrics: modularity and nestedness. Modularity measures the degree to which the network is divided in sets of nodes with high internal connectivity and low external connectivity (Olesen et al. 2007), which, in our case, identify plants and butterflies that interact more with each other than with other taxa in the network. Nestedness measures the degree to which the partners of poorly connected nodes form a subset of partners of highly connected nodes (Bascompte et al. 2003). To measure modularity, we used the Beckett (2016) algorithm, which works for both binary and weighted networks, as implemented in the function *computeModules* from the package *bipartite* (Dormann et al. 2008) in R version 3.6.2 (R Core Team 2019). This algorithm assigns plants and butterflies to modules and computes the modularity index, Q. To measure nestedness, we computed the nestedness metric based on overlap and decreasing fill, NODF (Almeida-Neto et al. 2008; Almeida-Neto and Ulrich 2011), as implemented for binary and weighted networks in the function *networklevel* also in the R package *bipartite*. To test when Q and NODF scores were significant, we computed standardized Z-scores that can be compared across networks of different sizes and complexities using the R package *vegan* (Oksanen et al. 2019) (details in Supplement).

We emphasize that our method does not estimate the first ages of origin for modularity or nestedness, but rather it estimates the first ages for which these network features can be detected. The difficulty of detecting network topological features increases with geological time, in part because phylogenetic reconstructions become less certain as age increases, but also because time-calibrated phylogenies of extant organisms are represented by fewer lineages as time rewinds. Ancestral network size and connectivity may therefore be underestimated. For these reasons, our statistical power to infer the age of origin for the oldest ecological interactions is limited. When interpreting our results, we focus on the ages that we first detect modularity and nestedness among surviving lineages, where first-detection times are assumed to postdate origination times for these network features.

Finally, we compared these estimates to the posterior distribution of *Z*-scores and statistical significance by calculating Q and NODF for 100 samples from the MCMC and 100 null networks for each sample. This comparison was done to test if the three summary networks accurately represent the posterior distribution of ancestral networks in terms of modularity and nestedness.

#### Posterior support for ecological modules

Defining eco-evolutionary groupings as modules allows us to visualize when those modules first appeared and how they changed over time. But in contrast to indices that are calculated for the entire network, the information about module configuration is not easily summarized into a posterior distribution. To circumvent this problem (and limitation 3 listed above), we used one of the weighted summary networks (probability threshold = 0.5) to characterize the modules across time, and then validate these modules with the posterior probability that two nodes belong to the same module (see below). This weighted network includes many more interactions than the binary network, while preventing very improbable interactions from implying spurious modules.

After identifying the modules for the summary network at each age, we assigned fixed identities to modules based on the host plant(s) with most interactions within the module. We then validated the modules in the eight summary networks (one for each time slice) using 100 networks sampled during MCMC, i.e. snapshots of character histories sampled at equal intervals during MCMC. We first decomposed each network of ancestral interactions sampled during MCMC into modules, and then calculated the frequency with which each pair of nodes in the summary network (butterflies and plants) were assigned to the same module across samples; that is, the posterior probability that two nodes belong to the same module.

### Bayesian inference method

We estimated the joint posterior distribution of model parameters and evolutionary histories using the inference strategy described in Braga et al. (2020) as implemented in the phylogenetics software, RevBayes (Höhna et al. 2016). Model parameters were assigned prior distributions of *μ* ~ Exp(10), *β* ~ Exp(1), and (λ_01_, λ_10_) ~ Dirichlet(1, 1). We ran four independent MCMC analyses, two with the anagenetic tree and two with the cladogenetic tree, each set to run for 2 × 10^5^ cycles, sampling parameters and node histories every 50 cycles, and discarding the first 2 × 10^4^ as burnin. To verify that MCMC analyses converged to the same posterior distribution, we applied the Gelman diagnostic (Gelman and Rubin 1992) as implemented in the R package *coda* (Plummer et al. 2006). Results from a single MCMC analysis are presented.

### Code availability

Our RevBayes and R scripts are available at https://github.com/maribraga/pieridae_hostrep. Our R scripts additionally depend on a suite of generalized R tools we designed for analyzing ancestral ecological network structures, which are available at https://github.com/maribraga/evolnets. An R package containing these tools is under development and will be officially released in a future publication.

## Results

Posterior estimates of Pieridae host-repertoire evolution were partially sensitive to whether we measured distances between host lineages in units of geological time or in units of major cladogenetic events (Fig. S2). When measuring anagenetic distances between host lineages, posterior mean (and 95% highest posterior density; HPD95) estimates were: global rate scaling factor for host-repertoire evolution *μ* = 0.02 (0.015 - 0.026), phylogenetic-distance power *β* = 2.1 (0.017 - 3.82), relative host gain rate λ_01_ = 0.035 (0.022 - 0.047), and relative host loss rate λ_10_ = 0.965 (0.95 - 0.98). Mean estimates were similar when distances between hosts were measured in units of cladogenetic events: *μ* = 0.019 (0.014 - 0.024), *β* = 1.48 (1.02 - 1.97), λ_01_ = 0.027 (0.017 - 0.036), and λ_10_ = 0.97 (0.96 - 0.98). Due to the stronger support for excluding *β* = 0 when using cladogenetic distances, we used the inferences from that analysis for the remaining results (see Fig. S3 for results with anagenetic distances). For the cladogenetic distance-based results, the posterior mean number of host-repertoire evolution events was 148 (75 gains, 73 losses) throughout the diversification of Pieridae. This is equivalent to approximately six events for every 100 million years of butterfly evolution (per lineage).

### Summarizing ecological interactions through time

Using the traditional approach for ancestral state reconstruction, that is, focusing on the highest-probability hosts at internal nodes of the butterfly tree, we described the general patterns of evolution of interactions between Pieridae butterflies and their host plants (Fig. 1). We could confidently say that: (1) the most recent common ancestor (MRCA) of all Pieridae butterflies used a Fabaceae host, (2) all ancestral Coliadinae and Dismorphiinae used Fabaceae, (3) the MRCA of, and early Pierinae (Pierina + Aporina + Anthocharidini + Teracolini) used a Capparaceae host, (4) Brassicaceae and Loranthaceae were each used by one Anthocharidini subclade, (5) early Aporina used both Loranthaceae and Santalaceae, and (6) the MRCA of Pierina and early descendants used three host lineages: Capparaceae, Brassicaceae and Tropaeolaceae.

**Figure 1.**
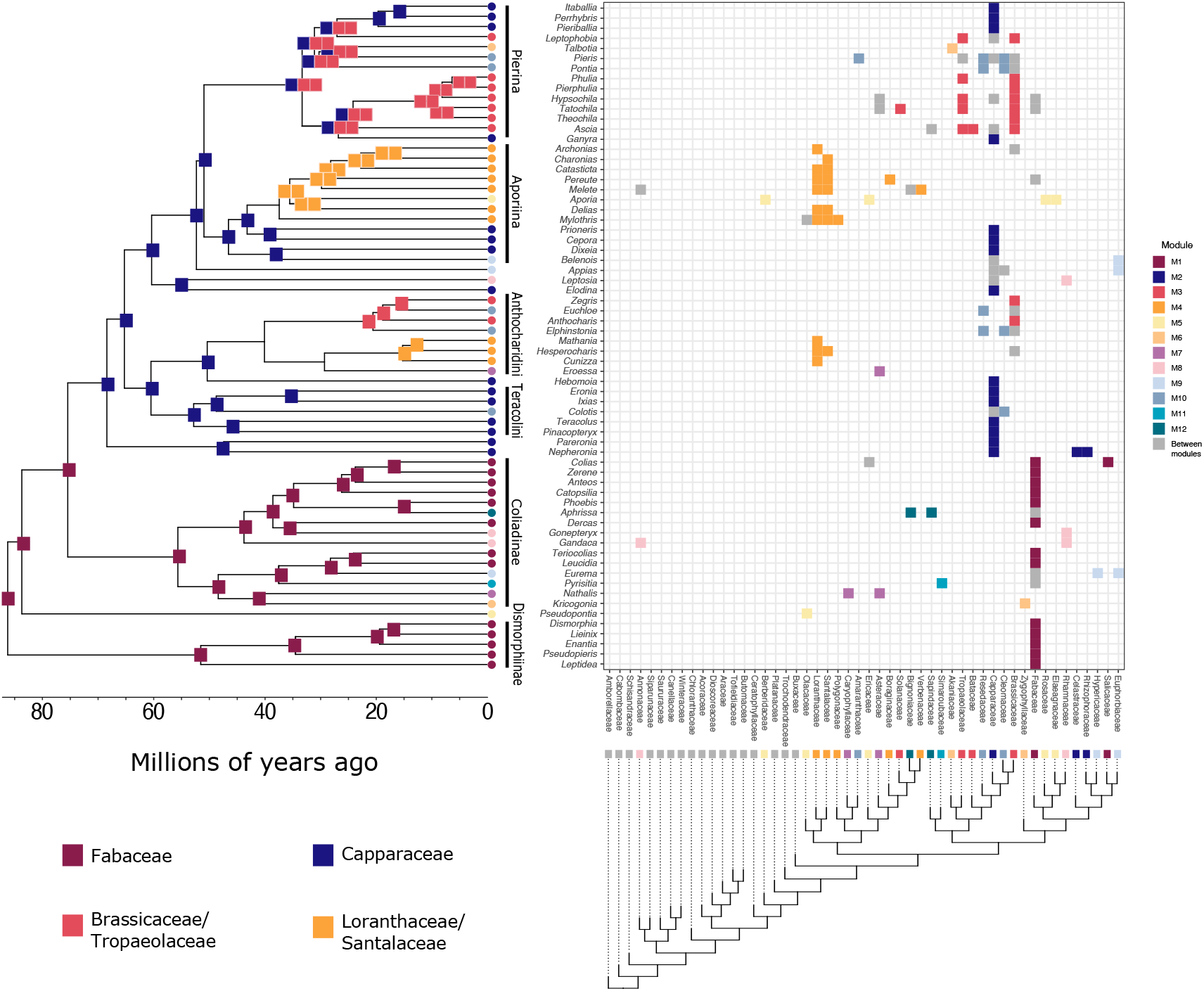
Ancestral state reconstruction showing interactions with marginal posterior probability ≥ 0.9. The model reconstructs how host repertoire evolved along the Pieridae phylogeny (left), based on the observed butterfly-plant interactions (top-right), and the cladogenetic distance between hosts (measured as the number of branches separating the hosts; bottom-right). The color of the symbols at the tips of both trees shows to which module the butterfly genus or plant family belongs (modules from the present-day network). Each square at internal nodes of the butterfly tree represents one plant family and is colored by the module to which the plant belongs. The matrix in the top-right shows the observed interactions between butterflies (rows) and plant families (columns). Rows and columns are ordered to match the phylogenetic trees. Interactions between butterflies and plants within modules are colored by module, whereas interactions between modules are in grey.

While the traditional ancestral state reconstruction described above reveals relevant and important pieces of the history of interaction between pierid butterflies and their host plants, it represents only a part of the posterior distribution of ancestral interactions. The remaining analyses provided more detailed information on the inferred host-repertoire evolution. Instead of reconstructing ancestral host repertoires at internal nodes of the butterfly tree, we looked at eight time slices along the diversification of Pieridae: every 10 Myr from 80 Ma to 10 Ma.

### Viewing ecological histories as networks

According to the posterior distribution of Q and NODF based on networks sampled from the MCMC, modularity and nestedness were first detectable 30 Ma (Fig. 2; for raw Q and NODF values see Fig. S4). But while the support for modularity has not changed much in the last 30 Myr, support for nestedness increased linearly in the past 50 Myr. Overall, the summary networks overestimated the presence of modularity, and only the weighted summary network with the 0.1-threshold correctly estimated that significant modularity first appeared at 30 Ma (Fig. 2 upper panel). On the other hand, the summary networks underestimated the existence of nestedness in ancestral networks (Fig. 2 lower panel), with several networks being significantly less nested than expected by chance, especially with the binary networks.

**Figure 2.**
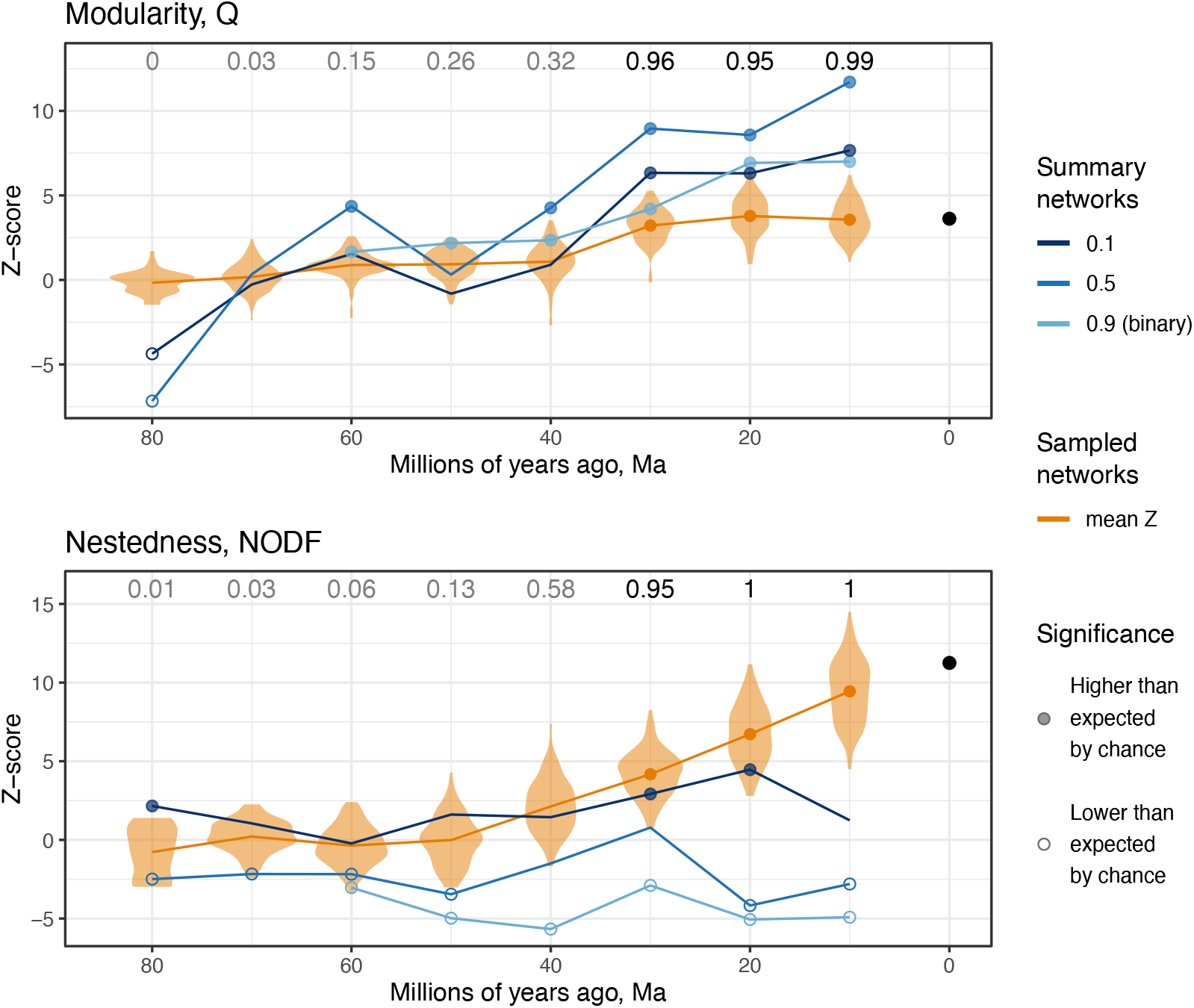
Structure of the Pieridae-angiosperm network over time. *Z*-scores for (a) modularity and (b) nestedness for summary (blues) and sampled networks (orange) from 80 Ma to 10 Ma, and for the observed present-day network (black circles). Each orange violin represents the distribution of *Z*-scores for sampled networks at each time slice and the orange line shows the mean *Z*-score. Indices (Q or NODF) higher than expected under the null model are shown with a filled circle, while indices lower than expected are shown with an empty circle. Numbers at the top of each graph show the proportion of sampled networks that were significantly modular or nested. In all cases, the significance level *α* = 0.05.

The present-day Pieridae-angiosperm network was characterized by both higher modularity (M = 0.64, p ≤0.001, *Z*-score = 3.62) and nestedness (NODF = 14.8, p ≤0.001, *Z*-score = 11.21) than expected by chance. Most of the butterfly lineages within Dismorphiinae and Coliadinae are associated with Fabaceae hosts (module M1), while Pierinae butterflies use many other host families (Fig. 1), the most common being Capparaceae (module M2), Brassicaceae + Tropaeolaceae (M3) and Loranthaceae + Santalaceae (M4). Interestingly, some Pierinae butterflies recolonized Fabaceae and others colonized new hosts while keeping the old host in their repertoire, resulting in among-module interactions that connected the whole network and produced signal for nestedness. By exploring the posterior distribution of ancestral interactions, we were able to characterize how this network was assembled throughout the diversification of Pieridae butterflies, as described below.

At 80 Ma, M1 and M2 were already recognized as separate modules in the summary networks (Fig. 3a). However, these modules were not validated by joint probabilities of two nodes being assigned to the same module across MCMC samples. Nodes that were assigned to different modules in the summary network were placed in the same module in many MCMC samples (grey cells in Fig. 4a). For example, Fabaceae and Capparaceae were assigned to the same module in 75 of the 100 MCMC samples, suggesting that at 80 Ma there was only one module including both Fabaceae and Capparaceae. Then, between the Late Cretaceous (represented by 70 Ma) and the Middle Eocene (represented by 50 Ma), Pieridae formed two distinct sets of ecological relationships with their angiosperm host plants: one set of pierid lineages feeding primarily on Fabaceae (M1), and a second set that first diversified between 70 and 60 Ma feeding primarily on Capparaceae (M2; Fig. 3b–d). It is important to note that even though we refer to this host lineage as Capparaceae throughout the network evolution, it likely diverged from Brassicaceae around 50 Ma (Magallón et al. 2015). Thus, before 50 Ma Capparaceae actually refers to the ancestor of Capparaceae + Brassicaceae. During that time, as more butterfly lineages accumulated within the Fabaceae and Capparaceae modules, the only plant lineages in the two modules were Fabaceae and Capparaceae themselves. Besides the two main modules, a small module formed around 50 Ma including the ancestor of *Pseudopontia* and Olacaceae.

**Figure 3.**
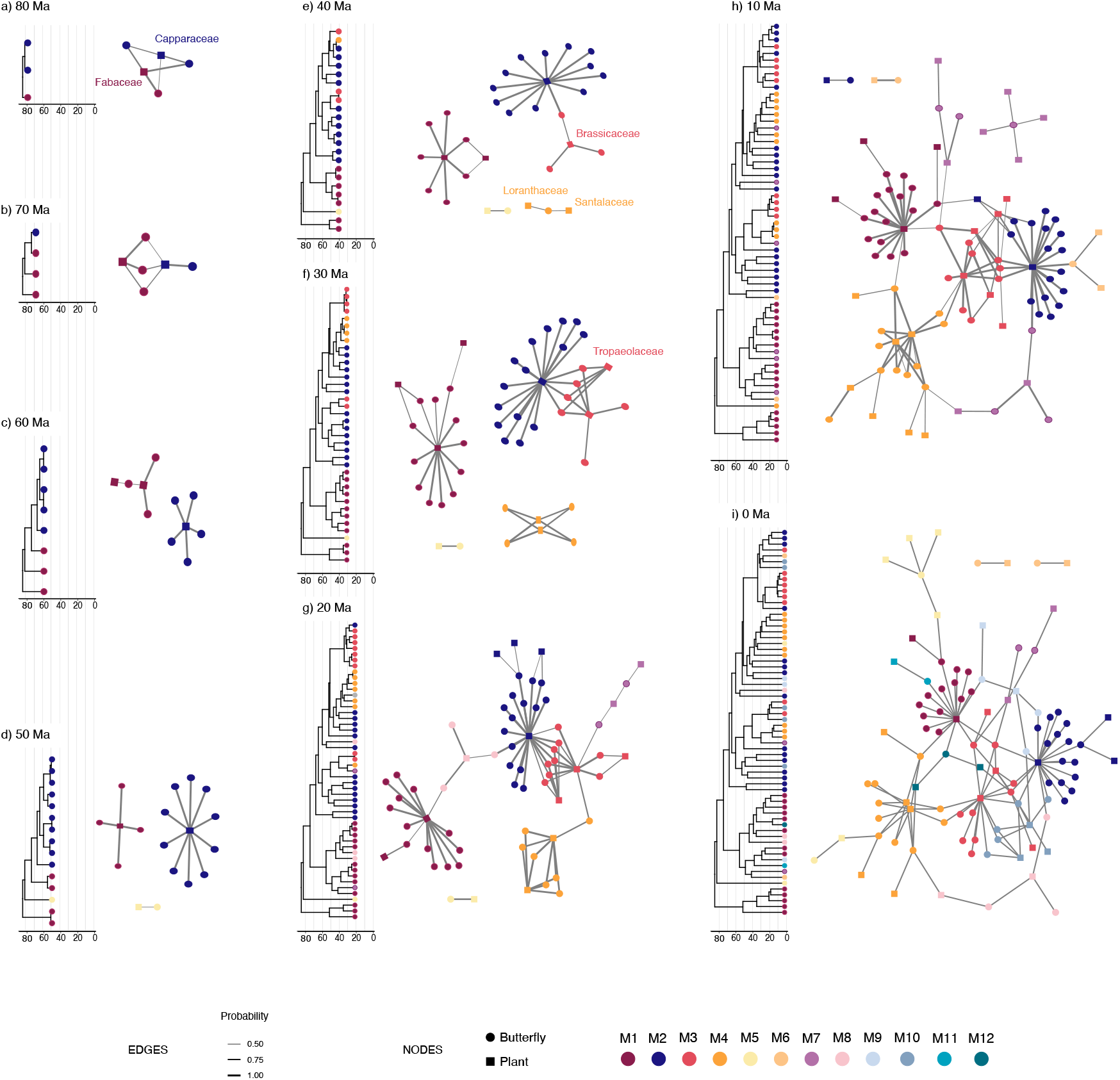
Evolution of the Pieridae-angiosperm network across nine time slices from 80 Ma to the present. Each panel (a-i) shows the butterfly lineages extant at a time slice (left) and the estimated network of interactions with at least 0.5 posterior probability (right). Edge width is proportional to interaction probability. Nodes of the network and tips of the trees are colored by module, which were identified for each network separately and then matched across networks using the main host plant as reference. Names of the six main host-plant families are shown at the time when they where first colonized by Pieridae.

**Figure 4.**
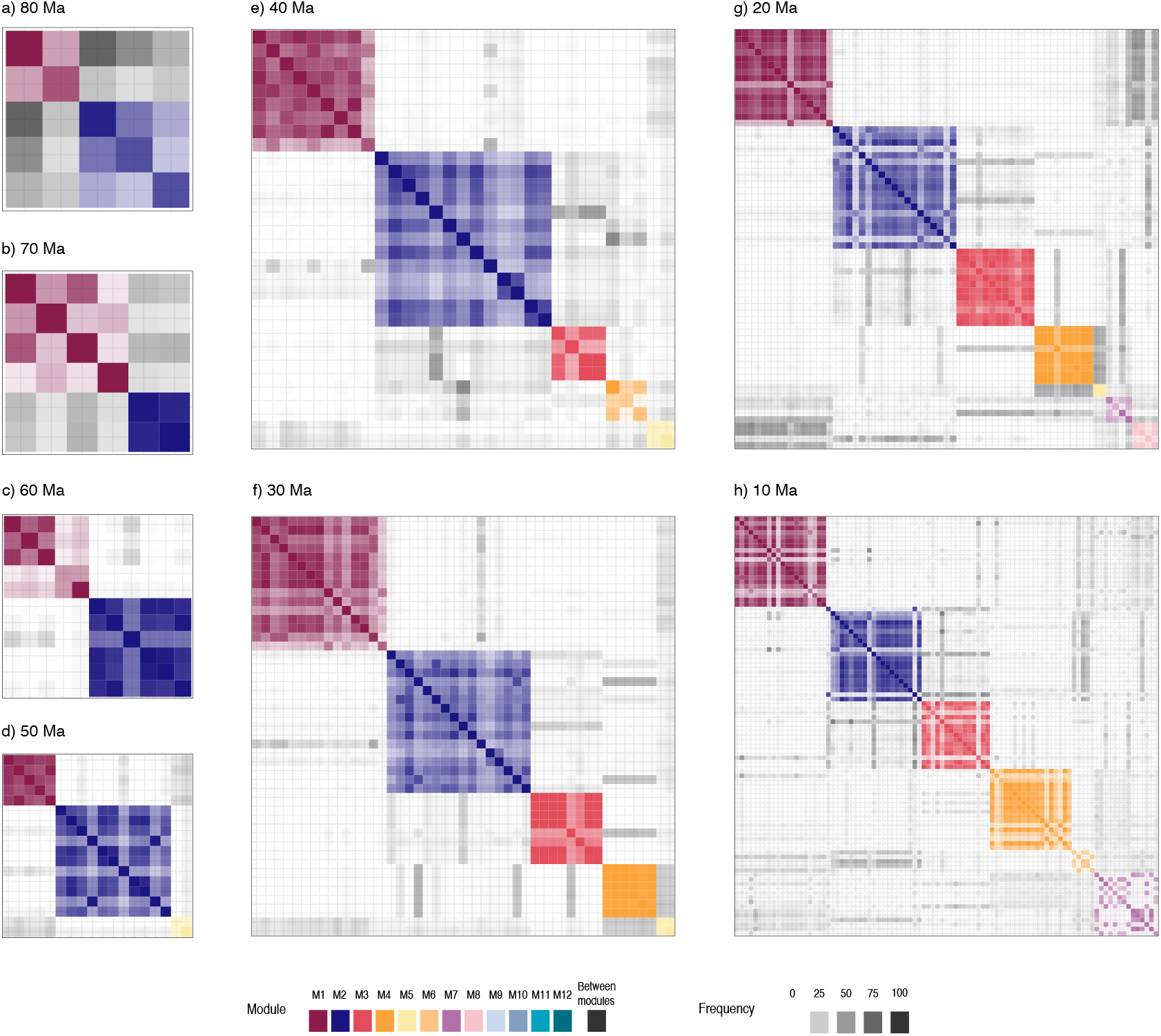
Heatmap of frequency with which each pair of nodes (butterflies and host plants) was assigned to the same module across 100 networks sampled throughout MCMC. In each panel, rows and columns contain all nodes included in the weighted summary network with probability threshold of 0.5 at the given time slice (depicted in Fig. 3). Rows and columns are ordered by module. When the nodes in the row and in the column are in the same module in the weighted summary network (Fig. 3), the cell takes the color of the module; otherwise, the cell is grey. The opacity of the cell is proportional to the posterior probability that the two nodes (row and column) belong to the same module.

Between 40 and 30 million years ago, coinciding with the onset of the Oligocene, two new modules emerged, one composed of butterflies that shared interactions with Brassicaceae and/or Tropaeolaceae (M3), and another of lineages that interacted with Loranthaceae and/or Santalaceae (M4; Figs. 4e–f and 5e–f). At the end of this period, M1 had expanded due to butterfly diversification and colonization of new host plants; M2 and M3 expanded and became more connected, as the first Pierina diversified while using both the ancestral host Capparaceae and the more recent host Brassicaceae. Entering into the Miocene at 20 Ma and 10 Ma, as the sizes of modules grew, so did the number of interactions between modules. Modules M6, M7 and M8 appeared for the first time, and the remaining modules, M7–M12, appeared between 10 Ma and the present.

## Discussion

We developed a novel methodological framework to reconstruct ancestral networks of ecological interactions between two clades from posterior distributions of evolving host repertoires. Our approach allows researchers to characterize ancestral ecological networks while accounting for uncertainty in ancestral state estimates, to measure the probability that any species-pair was assigned to the same ancestral module, and to identify when network structures (modularity and nestedness) first became statistically detectable in evolutionary time. Applying our framework to the Pieridae-angiosperm system, we tested the ideas proposed in Braga et al. (2018), who suggested that the evolution of butterfly-plant interactions was shaped by a combination of processes consistent with both the escape-and-radiate (Ehrlich and Raven 1964) and oscillation hypotheses (Janz and Nylin 2008), and inferred that certain evolutionary changes in host use would leave recognizable features in ecological networks. We found that seven (out of 75) host gains of five plant families had an outsized effect on Pieridae-angiosperm network structure, creating and connecting the main modules: Capparaceae (gained once), Loranthaceae (twice), Santalaceae (once), Brassicaceae (twice), and Tropaeolaceae (once). We discuss our empirical findings below, focusing on how three types of evolutionary change in host use restructured ancestral Pieridae-angiosperm networks.

First, a complete host shift (i.e. gain of new host followed by loss of ancestral host) can produce a new module isolated from the rest of the network. Our reconstructions are consistent with the hypothesis that early Pierinae butterflies shifted hosts from Fabaceae (Fabales) to Capparaceae (Brassicales), resulting in the creation of a new ecological module in the network (M2 in Figs. 2, 4 and 5). The diversification of Pierinae was first explained as a radiation following the colonization of the chemically well-defended Brassicales plants (Ehrlich and Raven 1964; Braby and Trueman 2006). More recent studies identified the origins of defense and counter-defense mechanisms, supporting the idea of an arms-race during Pierinae-Brassicales coevolution (Wheat et al. 2007; Edger et al. 2015). All evidence from the present and the aforementioned studies suggest that the host shift from Fabaceae to Capparaceae completed between 70 and 60 Ma. The timing of this shift overlaps with the Cretaceous—Paleogene (K-Pg) extinction event and aligns with a previously estimated increase in Brassicales diversification rate (Edger et al. 2015). Although we cannot conclude if and how the K-Pg extinction event and a coevolutionary arms-race induced a shift in host use by Pieridae, these factors were likely involved in the origin of the Pierinae-Brassicales association.

Two other important host shift events occurred in the Late Eocene and Early Oligocene (ca. 30-40 Ma) that contributed to the formation of modules M3 and M4. Early ancestors of Anthocharidini (the sister clade to the Capparaceae-using *Hebomoia*) shifted from Capparaceae to the early Brassicaceae and joined with Pierina to form module M3 (discussed below). At a similar time, as Aporiina first diversified, one of its earliest lineages shifted hosts from Capparaceae to instead use the closely related Loranthaceae and Santalaceae, thus creating module M4. Between the Late Eocene and the present, the number of Pieridae genera associated with the newly-formed M3 and M4 modules grew to rival the number of genera belonging to modules M1 and M2.

In the second type of host-use change, host-range expansion (i.e. colonization of new hosts without loss of ancestral host) increases the size of the module and creates nestedness within the module. Our method, which notably allows butterflies to simultaneously use multiple hosts, inferred that early Pierina lineages (ca. 35 Ma) used three plant families: Capparaceae and – the two newly acquired – Brassicaceae and Tropaeolaceae (Fig. 1). More recently, the two recent ancestors of *Archonias* and *Hesperocheris* independently expanded their host repertoires from Loranthaceae/Santalaceae alone (M4) to include Brassicaceae (M3) in the past 15 to 20 Ma. These host-range expansions, which helped form and interconnect module M3, coincided with the origin of the Core Brassicaceae and increases in diversification rates in both Pierina and Brassicaceae (Edger et al. 2015), thus having a major effect on network dynamics by increasing network size and nested substructures.

Third, recolonization (i.e. gain of a host that has been used in the past) connects different modules and increases global nestedness. One (or possibly two) Pierina lineages that were historically associated with Brassicaceae (M3) recolonized Fabaceae (M1) in the Early Miocene (ca. 20 Ma), connecting the newer M3 module with the M1 module. By 10 Ma, ancestors of *Pereute* in the Aporiina clade, which primarily uses and used Loranthaceae/Santalaceae hosts (M4), united module M4 with M1 by also recolonizing an ancestral host, Fabaceae. In both lineages, Fabaceae was recolonized for the first time in roughly 40 Myr, since 65 Ma, at least. After modules M3 and M4 connected to module M1 through recolonization, modules M1 and M4 gained new indirect connections to Capparaceae (M2) through Brassicaceae (M3).

The three event types we discussed above had an aggregate effect upon how global properties of network structure evolved over time. Modularity and nestedness were first detected between 40 Ma and 30 Ma (Fig. 2). While network size increased in the last 30 Myr, most interactions occurred within each of the four largest modules (M1–M4). Most likely, modularity increased with establishment of the M3 and M4 modules, while nestedness emerged because early Pierina retained Capparaceae as a host in its repertoire, thereby connecting modules M2 and M3. While modularity remained almost constant in the past 30 Myr, nestedness increased linearly over time (Fig. 2). Seven modules that were first detected in the past 30 Myr were connected to at least one, but often two, of larger modules. In effect, as butterflies gained new hosts and formed new modules, a subset of these butterflies retained or recolonized their ancestral hosts (Fabaceae, Capparaceae, Brassicaceae, Tropaeolaceae, Loranthaceae or Santalaceae, depending on the butterfly clade), preserving connectivity to the original modules. Thus, host-range expansions and recolonizations promoted a phase transition in the basic structure of the network (Guimarães Jr. 2020), which went from a disconnected network composed of small, isolated modules to an interconnected network with a giant component (Newman et al. 2001). This is an important example for how giant components emerge in ecological networks, which allow eco-evolutionary feedbacks to propagate across multiple species in an ecosystem.

Biological descriptions, rather than methodological explanations, for how specific modules evolved lend credibility to our inferences as a whole. For example, module M3 resulted from the colonization of Brassicaceae, which is closely-related to Capparaceae, but better chemically defended. Shortly after the appearance of Brassicaceae, two lineages within Pierinae colonized the family and subsequently diversified, probably by evolving counter-defense mechanisms (Edger et al. 2015). Tropaeolaceae, also in module M3, was colonized by the Pierina, but these plants have no indolic glucosinolates as chemical defense. When colonization occurred is less clear. While some suggest that Pierina and Tropaeolaceae first interacted in the Holocene, based on their historical distribution (Edger et al. 2015), our analysis suggests a much older colonization. One possible explanation is that Pierinae gained the ability to use Tropaeolaceae (and possibly other Brassicales lineages) when they first colonized Brassicaceae, but the interaction was only realized when their geographical distributions overlapped. Module M4, on the other hand, was likely formed by the colonization of hemiparasitic plants (Santalaceae and Loranthaceae) growing on Brassicales hosts (Braby and Trueman 2006).

Our key findings depend on our ability to reconstruct the evolution of ecological networks accurately. Existing methods designed for fast-evolving bacteria-phage systems (Weitz et al. 2013), simulation-based pattern assessments (Guimarães et al.2017; Braga et al. 2018), and tanglegram-based methods (de Vienne 2018), though suitable for many problems, could not be used to reconstruct how timed sequences of ancestral host gain and loss events induced structural changes to ecological networks over deep time scales. Researchers who are interested in investigating coevolutionary problems similar to those we examined in the Pieridae-angiosperm system will also find our method useful. Future versions of the method can incorporate additional forms of biological evidence, including traits (e,g. chemical attractants or deterrents; Levin 1976; Ramírez et al. 2011), biogeography (Hoberg and Brooks 2008; Hembry et al. 2013), fossils (Opler 1973), and a wider range of mutualistic and/or antagonistic interactions (Bascompte and Jordano 2007; Satler et al. 2019; Wang et al. 2019). Several statistical properties of the method also deserve further attention in subsequent research, including how to accommodate phylogenetic uncertainty (e.g. by using credible sets of butterfly and plant phylogenies), how to account for the potential influence of extinct or unsampled lineages upon network reconstruction, how to define and compute consensus modules across posterior samples, and how to assign biologically meaningful module labels across posterior samples and time periods.

In summary, the diversification and evolution of host repertoire of Pieridae butterflies can indeed be explained by a combination of the escape-and-radiate (Ehrlich and Raven 1964) and the oscillation hypothesis (Janz and Nylin 2008). Although others have also suggested that both hypotheses are at play (Segar et al. 2017; Braga et al.2018), here we provide evidence for the mechanistic basis of host-repertoire evolution that underlies network evolution. Our results demonstrate the power of combining network analysis with Bayesian inference of host-repertoire evolution in understanding how complex species interactions change over time. Future avenues of research should explore the extent to which host shifts, host-range expansions, and host recolonizations characterize the evolution of other networks of intimate interactions. Given the support for the oscillation hypothesis from a variety of systems such as polyphagous moths (Wang et al. 2017), parasites (Hoberg and Brooks 2008; D’Bastiani et al. 2020), and even plant-microbial mutualism (Torres-Martínez et al. 2021), we expect similar dynamics to be found in many systems, ranging from parasitisms to mutualisms.

## Acknowledgements

We thank Paulo R. Guimarães Jr. and Christopher W. Wheat for discussing and commenting on earlier versions of this paper, and four anonymous reviewers provided valuable feedback that also improved the clarity and focus of the manuscript. The Norwegian Institute for Nature Research in Tromsø kindly provided workspace and research infrastructure to M.P.B. during manuscript preparation. The Swedish Research Council supported [2015-04218 to S.N.] and [2014-05901 to F.R.]. M.J.L. was supported by startup funds from WUSTL.

## Statement of authorship

MPB, NJ and SN designed the basis for the biological study. SN collected the data. MPB and MJL designed the statistical analyses. MPB analyzed the data, generated the figures, and wrote the first draft of the manuscript. All authors contributed to the final draft.

## Data accessibility statement

No new data were used.

## Notes

https://github.com/maribraga/pieridae_hostrep

### Competing Interest Statement

The authors have declared no competing interest.

### Summary of Updates

The title was updated; Figure 1 was moved to the Supplementary; Introduction and Discussion were updated.

