## Supplementary for "Phylogenetic reconstruction of ancestral ecological networks through time for pierid butterflies and their host plants"

Version dated: June 16, 2021

**- Reference list used to gather host use records for Pieridae butterflies**

**- Supplementary methods: Z-score**

**- Figure S1: Time-calibrated phylogeny of Pieridae.**

**- Figure S2: Posterior densities for model parameters.**

**- Figure S3: Ancestral state reconstruction of Pieridae host repertoire.**

**- Figure S4: Raw modularity and nestedness across the evolution of the Pieridae-angiosperm network.**

*

**Supplementary methods: Z-score**

We investigated whether changes in modularity and nestedness could be detected over evolutionary time by computing $Z$-scores for Q and NODF statistics for the observed extant network and for each of the three summary networks at each age. $Z$-scores and statistical significance for modularity and nestedness were determined by comparing the observed values of Q and NODF to those of a null distribution. Each null distribution was generated using the *nullmodel* function from the R package *vegan* (Oksanen et al. 2019) to simulate 1000 null networks. Each null network had the same number of nodes and edges as the observed network (but not the same distribution of edges per node), and edges between nodes were sampled uniformly at random. All Q and NODF values were standardized as $Z = \frac{x-\mu}{\sigma}$, where $x$ is the Q or NODF statistic that is observed in a particular network, $\mu$ is the mean value and $\sigma$ is the standard deviation for the statistic under the null distribution. This standardisation quantifies the position of the observed metric within the null distribution in standard units (Ulrich et al. 2009), allowing us to compare different networks.

*

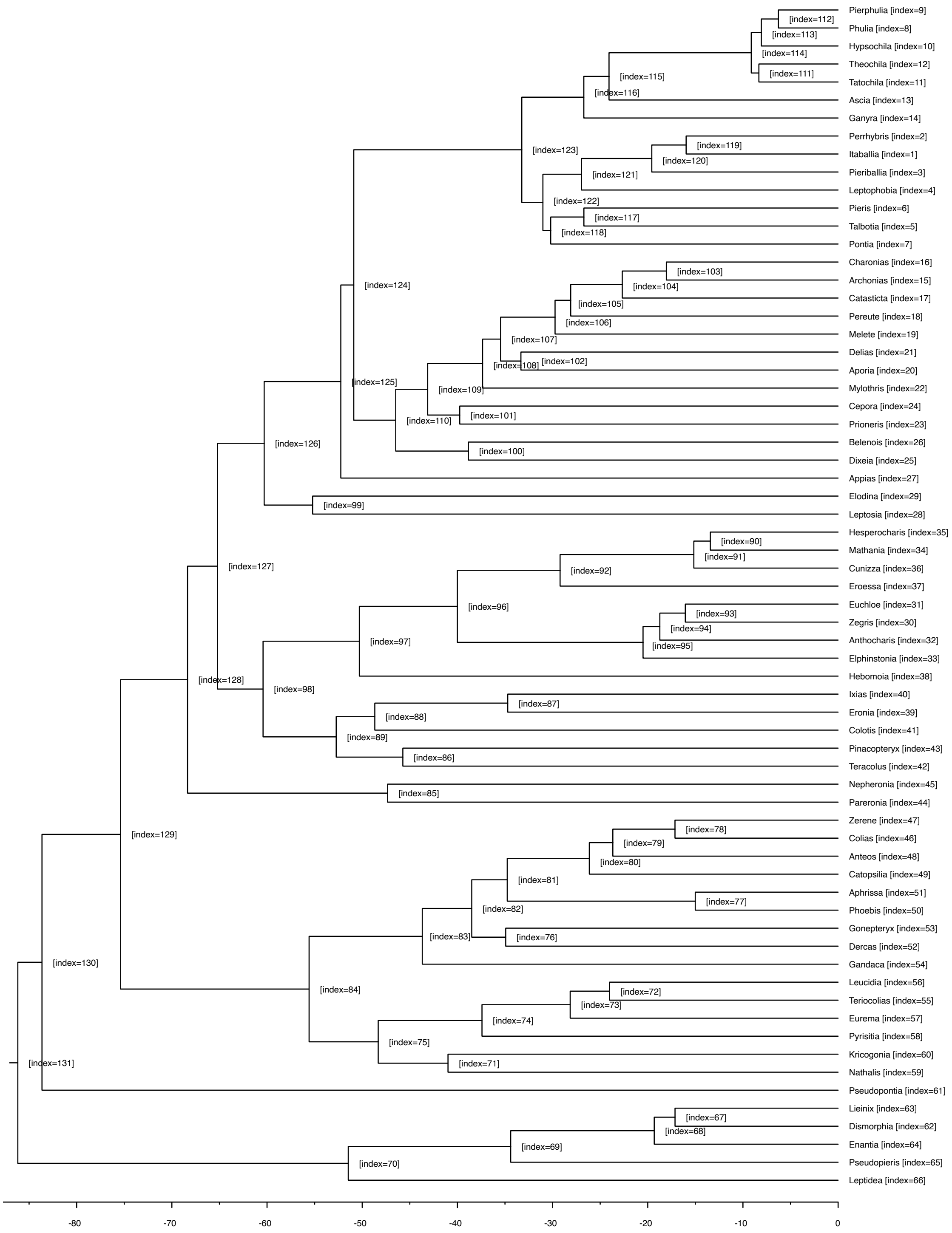


Figure 1: Time-calibrated phylogeny of Pieridae butterflies used in this study. Indices of internal nodes correspond to row names in Figure S2.

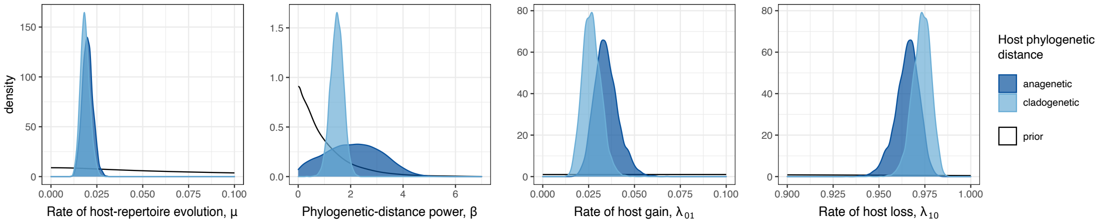


Figure 2: Estimated marginal posterior densities for parameters in the host-repertoire evolution model using two different representations of the phylogenetic distance between host-plant families: anagenetic (time) or cladogenetic (number of branches).

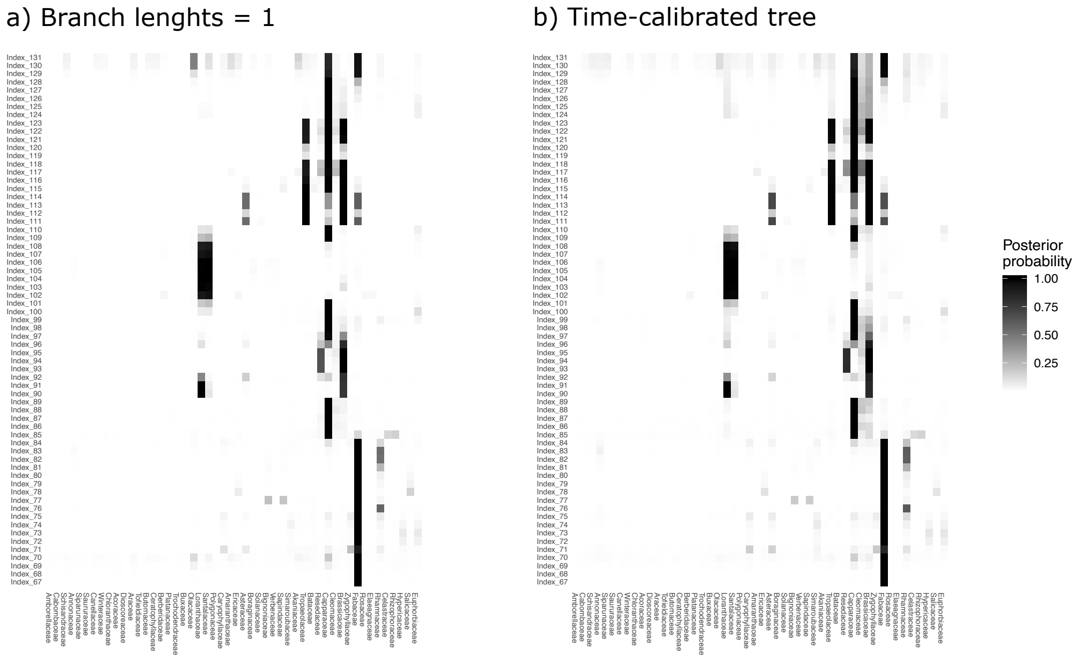


Figure 3: Posterior probability of ancestral host repertoires of Pieridae butterflies for three different model settings: a) host tree with branch lengths assigned to 1; b) host tree with branch lengths proportional to divergence time; and c) independence model ($\beta = 0$). In each panel, rows show the host repertoires at internal nodes of the Pieridae phylogeny (same index as in Fig. S1). The probability of each host being on the host repertoire of each butterfly is shown by the color scale.

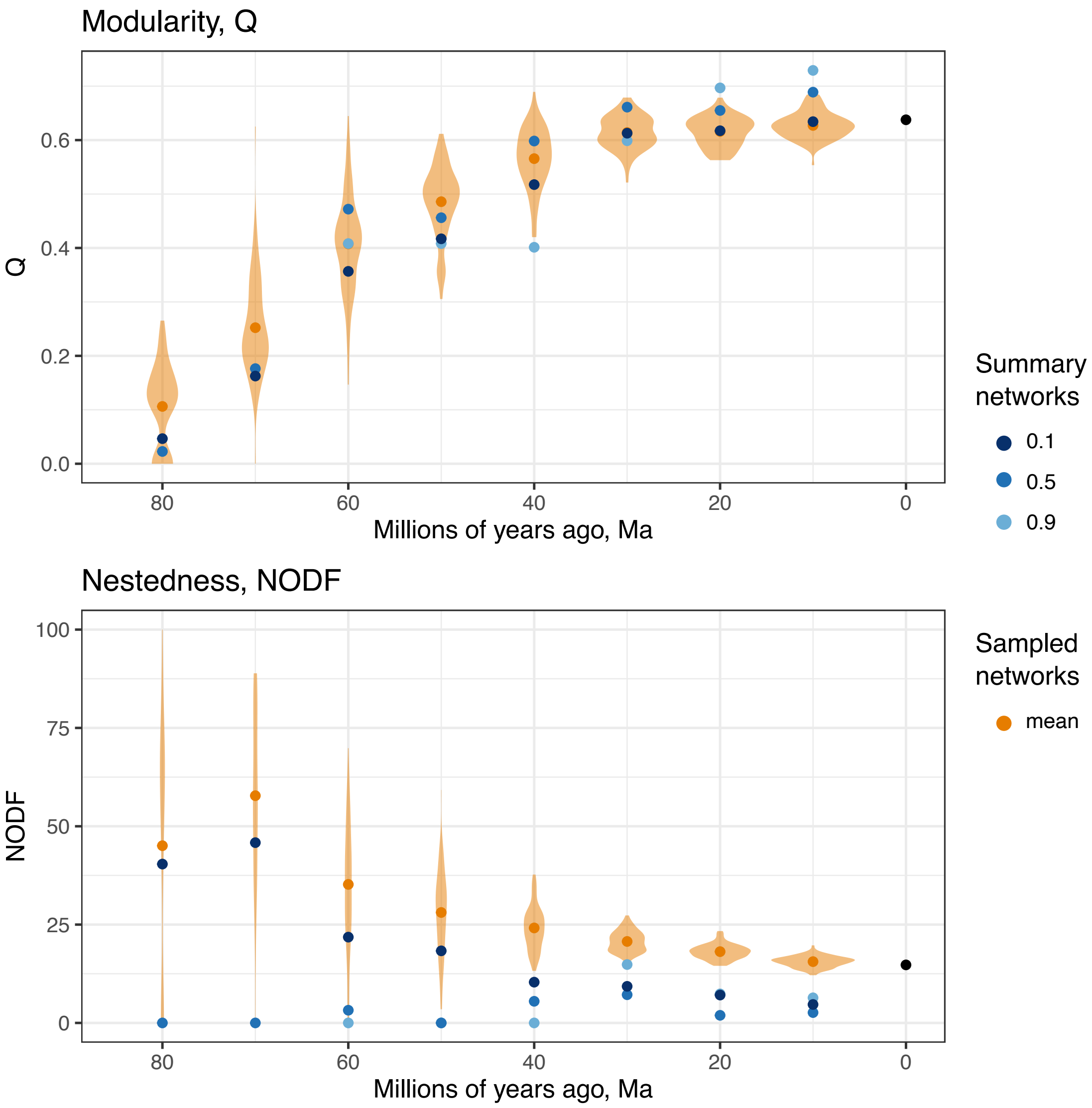


Figure 4: Modularity and nestedness for summary (blues) and sampled networks (orange) from 80 Ma to 10 Ma, and for the observed present-day network (black circles). Each orange violin represents the distribution of the index (Q or NODF) for sampled networks at each time slice and the orange circle shows the mean.
